# Emergence of reward expectation signals in identified dopamine neurons

**DOI:** 10.1101/238881

**Authors:** Luke T. Coddington, Joshua T. Dudman

## Abstract

Coherent control of purposive actions emerges from the coordination of multiple brain circuits during learning. Dissociable brain circuits and cell-types are thought to preferentially participate in distinct learning mechanisms. For example, the activity of midbrain dopamine (mDA) neurons is proposed to primarily, or even exclusively, reflect reward prediction error signals in well-trained animals. To study the specific contribution of individual circuits requires observing changes before tight functional coordination is achieved. However, little is known about the detailed timing of the emergence of reward-related representations in dopaminergic neurons. Here we recorded activity of identified dopaminergic neurons as naïve mice learned a novel stimulus-reward association. We found that at early stages of learning mDA neuron activity reflected both external (sensory) and internal (action initiation) causes of reward expectation. The increasingly precise correlation of action initiation with sensory stimuli rather than an evaluation of outcomes governed mDA neuron activity. Thus, our data demonstrate that mDA neuron activity early in learning does not reflect errors, but is more akin to a Hebbian learning signal - providing new insight into a critical computation in a highly conserved, essential learning circuit.

## Main text

Whether reaching to grab a morsel of cheese or to press an elevator button, the initiation of purposive action is associated with an expectation about its outcome. There are two broad conceptualizations of how learning about outcomes occurs in the brain. The first is a gradual strengthening of an association between an action or a stimulus and a specific outcome – often described as Hebbian learning^1^. The second is a gradual reduction in errors through comparison of expected and observed outcomes - often invoked to explain reinforcement learning^2^. It has been suggested that dissociable brain circuits and cell-types preferentially participate in one or the other learning mechanism^3^. For example, there is substantial evidence that the activity of midbrain dopamine (mDA) neurons primarily or even exclusively reflects error signals in well-trained animals^4^. Learning in a novel reward context can, however, differ fundamentally from the adaptation of steady state behavior in a familiar context. Consider, for example, the difference between an expert guitarist learning a new melody and a novice attempting to play the same melody for the first time. Whereas the trained guitarist will make meaningful errors that can be used to refine performance, the novice might only learn from the occasional successfully executed note.

Although it is clear that mDA neuron activity correlates with errors in the prediction of rewards following extensive training^4^, the circuit mechanism by which errors are computed remains ambiguous^5^. An important open question remains: as an animal begins to learn about rewards and informative sensory stimuli in its environment how do expectations about rewards first form? Does the ability to represent errors form at the same time as an expectation, or perhaps only after a robust, relatively accurate expectation is formed? Currently, there is no data examining the continuous emergence of phasic responses in the activity of single, identified mDA neurons as animals transition from a naïve state through learning. Even in the cases where measurements of mesolimbic dopamine release have been made across learning^6,7^ only a correlation with steady-state behavior has been established.

The neural correlates of initial, naïve learning have been impractical to ascertain for several reasons. First, traditional electrophysiological techniques make it difficult to unambiguously identify dopamine neurons, especially if their responses are non-canonical. Second, the earliest detectable changes in activity during initial learning may be subtle, necessitating very high signal to noise recordings of activity. Third, the difficulties in targeting a deep midbrain structure for recording and the brief, unique windows of novel learning for each subject hurt the yield of recorded cells. We addressed these issues by making juxtacellular recordings from optogenetically-identified (‘optotagged’) mDA neurons in the ventral tegmental area (VTA, n =47) and substantia nigra pars compacta (SNc, n =88) of head-fixed, thirsty, naïve mice as they learned an auditory trace conditioning paradigm (Figure 1; Methods). Localization of the SNc or VTA on each penetration was determined by observing the magnitude of an optogenetically-evoked field potential (Fig. 1c-e) known to reflect the presence of mDA neurons^8^. Juxtacellular recordings were obtained in the region of the largest evoked potential and mDA neuron identity was confirmed by driving trains of action potentials (Fig. 1e). The range of depths over which evoked potentials and positively identified mDA neurons were found agreed well with the anatomy of the VTA and SNc (Fig. 1f).

**Fig. 1.**
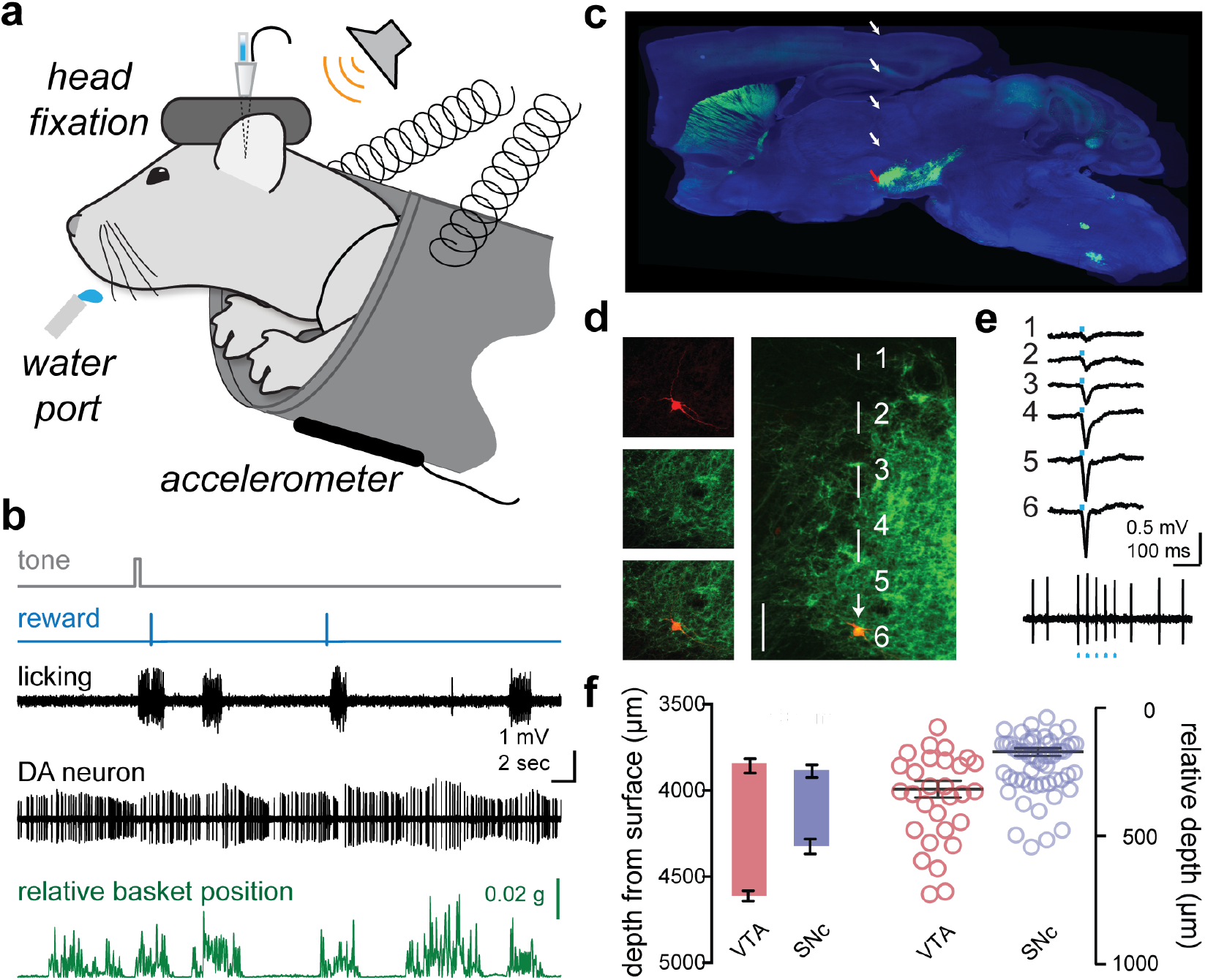
Juxtacellular recording from identified midbrain dopamine neurons in awake, behaving mice. **a)** Schematic of the head-fixed behavioral apparatus. Mice were head-fixed in front of a water reward port and supported by a spring-suspended basket monitored by an accelerometer. **b)** Auditory stimuli, openings of a solenoid water valve (reward), vibration of the water port due to licking (licking), and juxtacellular recordings from identified midbrain dopamine (mDA) neurons could be recorded simultaneously with bodily movements that displaced basket position relative to its resting position (relative basket position). **c)** mDA neurons in DAT-cre x ai32 mice selectively express a channelrhodopsin-YFP (ChR2-YFP) fusion protein. Representative histological section shows robust labeling of mDA soma in the midbrain and terminals in the striatum. White and red arrows indicate the trajectory of a recording pipette and approximate location of recorded neuron for the example experiment shown in D-E. **d)** (left) Individual and merged images of neurobiotin-labeled neuron (red) and YFP-expressing mDA neurons (green). (right) Pipette trajectory with locations of light stimulations numbered. **e)** Example traces of light-evoked field potentials corresponding to the recording locations (1-6) shown in D, demonstrating tight correspondence between ChR2-YFP expression (green) and evoked field potential amplitude. Lower trace shows resulting juxtacellular recording with tight entrainment of action potentials to the light stimulus (optogenetic tagging). **f)** Spatial distribution of both VTA (red) and SNc (blue) optogenetically tagged mDA neurons either relative to the dorsal surface of the brain (left) or relative to the dorsal-most position at which a light-evoked field potential was observed.

A further practical challenge is the inference of learning from the characteristically unreliable behavioral output of novice animals. While head-fixation reduced much potential variability, we also measured two independent behavioral responses to expectation of a water reward: licking at the reward spout and preparatory movement of the body. To quantify body movement, we supported mice in a spring-suspended basket equipped with an accelerometer (Fig.1a-b, see Methods). Note that body movements were correlated with, but not synonymous with, bouts of licking (Fig. 1b) – indicating that body movements can provide an additional metric to track learned behavior.

In our trace conditioning paradigm a 0.5 sec long auditory tone was presented 1.5 sec before delivery of a water reward (Fig. 1a-b; see Methods). Learning that the tone predicted water delivery was evidenced by steady increases in anticipatory behavior, as quantified by the number of licks during the delay between tone presentation and water delivery, as well as an increase in attempted body movement towards the water port (Fig. 2a-b). Whereas mice overtrained on this same paradigm rapidly re-acquire and extinguish responses to the tone^9^, naïve learning is predictably characterized by a much slower emergence of learned behavior (Fig. 2a). Changes in behavior associated with learning first emerged as isolated licks and small movements shortly following the tone onset (Fig. 2a, sessions 1-3, “early” acquisition), then developed into sustained lick bouts combined with large, sustained body movements (Fig. 2a, sessions 4-8, “middle” acquisition). While robust body movements initially followed water delivery, these movements were gradually attenuated late in training—reflecting a shift of movement initiation to precede water delivery (Fig. 2a, sessions 9+, “late” acquisition). The amount of licking and movements following the tone onset changed monotonically with training trials (relative difference in basket displacement in response to the tone and reward, r = −0.43, p < 0.0001; latency to initiate licking following tone, r = −0.50, p < 0.0001; licks in the CR during tone-reward interval, r = 0.63, p < 0.0001, Fig. 2b).

**Fig. 2.**
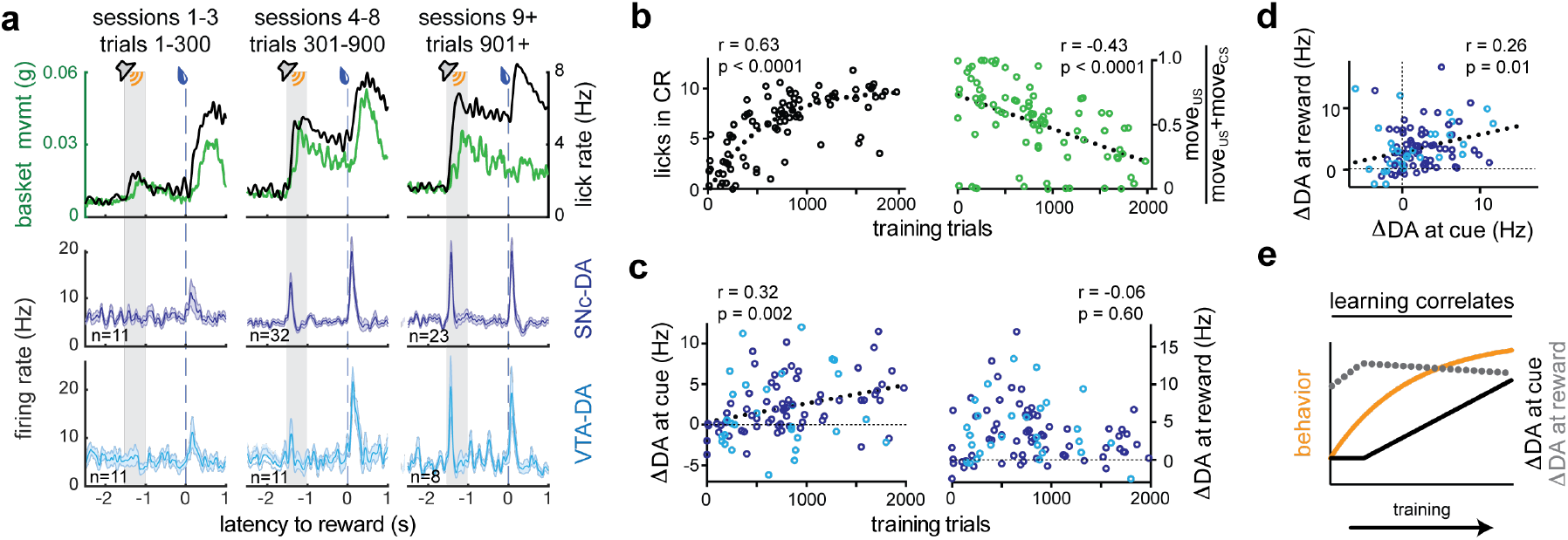
mDA neuron responses to predictive cue and reward stimuli evolve independently during acquisition learning. **a)** Behavioral (upper row) and physiological (middle and lower rows) data aligned to the opening of the water valve and divided in to early (sessions 1-3, left column) middle (sessions 4-8, middle) and late (sessions >8, right) training sessions. Conditioned responding was assessed by measuring relative basket position (basket mvmt) and lick rate. Smoothed peri-event time histograms (PETHs) for identified mDA neuron populations recorded in SNc (purple, middle row) and VTA (cyan, lower row). Shaded area indicates the standard error of the mean (s.e.m.) number of cells composing the mean is labeled for each panel. **b)** Behavioral learning measures during each individual neuron recording period. Both the number of licks during the cue-reward trace interval (licks in CR, left) and basket movement (fraction of movement following the reward) were significantly correlated with the number of training trials experienced at the time of recording. **c)** The mean modulation of mDA neuron activity following the predictive tone (left) or reward (right) were plotted as a function of the number of training trials experienced at the time of recording. Trend lines are indicated with thick dashes, y=0 highlighted as thin dashed lines. Colors reflect SNc (purple) and VTA (cyan) populations. **d)** Mean modulations of mDA neurons in response to predictive tone and reward stimuli are positively correlated **e)** Abstract summary of correlations of data from (a)-(d) across training

SNc and VTA mDA neuron populations lacked a coherent response to the tone early in learning, but developed robust phasic responses to the reward predictive tone as training progressed (r = 0.32, p = 0.002, n = 96, Fig. 2a, c). To our surprise we found that the proportion of mDA neurons responding to water delivery *also* increased over the first three sessions (1-3 sessions: 10 of 22, 4-8 sessions: 38 of 43, 9+ sessions: 27 of 31), while individual mDA response amplitude was not correlated with the number of trials experienced (r = −0.06, p=0.6, n = 96, Fig. 2c). The inference from existing data examining trained animals was that mDA neuron activity should transfer from a reward to an earlier, predictive stimulus across learning^4,6,7,10,11^. By contrast, our data – the first to examine the time course of the emerging mDA response to a predictive stimulus in detail – suggest that mDA responses to the predictive stimulus and the reward might be independent. One concern is that a transfer from reward delivery to an earlier predictive stimulus within individual neurons could have been obscured by population variability. Thus, we examined the correlation of tone and water delivery responses within individual mDA neurons. However, we observed that tone and water delivery responses were *positively* correlated in individual mDA neurons (r = 0.26, p = 0.01, n = 96; Fig. 2d). Our data are thus consistent with mDA responses that develop independently to the tone and water delivery but are scaled by a common cell-specific factor (Fig. 2e; Supplementary Fig. 1)^12,13^.

Correlation between the mDA neuron responses to reward-predictive tones and rewards themselves implies that these responses are representations of a common variable. We note that even in primary thirst and hunger centers in the rodent brain, modulation of activity in fact precedes consumption and so can be predictive of the reward available rather than strictly evaluative of the reward received^14,15^. Thus, we next asked how the mDA neuron response to reward delivery emerged during the earliest stage of learning. We found that modulation of mDA excitation within the first 3 training sessions in fact preceded contact of the tongue with the water port (Fig. 3). Note, this is at a point in training before any detectable mDA responses to the tone, and as training progresses reward responses evolve to even further lead reward receipt (Fig. 3c) as mice became familiar with the sensory stimuli attending reward delivery. These results were repeated in separate mice trained using an inaudible water delivery system (Fig. 3d), ruling out a role for a salient auditory cue associated with water delivery. This suggests that the dopamine response to reward delivery reflects the initiation of appetitive behavior – a bout of consummatory licking – rather than an evaluative response attending consumption of the reward. Thus, the correlated mDA responses to a reward-predictive tone and water delivery could reflect the fact that both events reflect reward expectation.

**Fig. 3.**
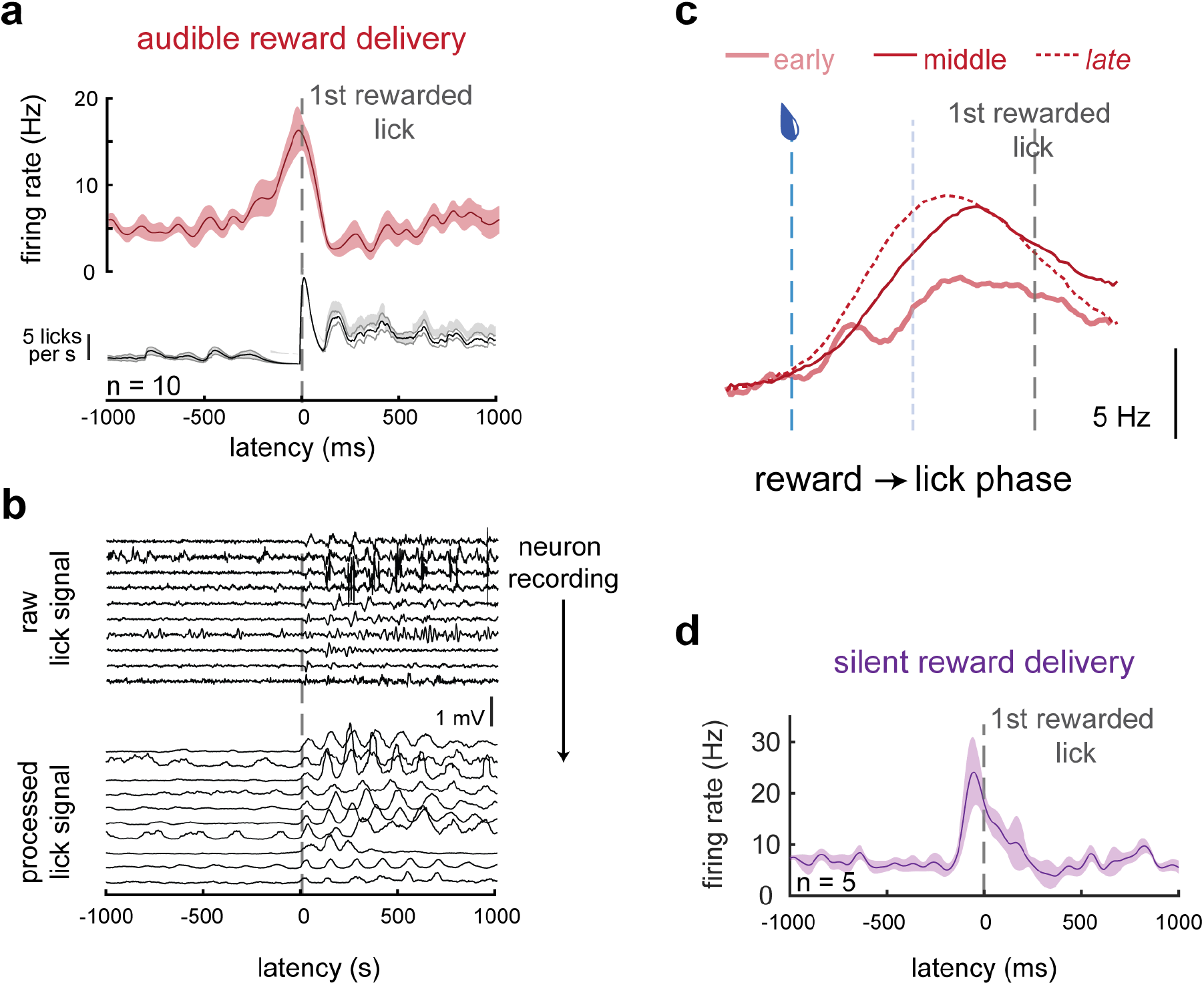
mDA reward signaling is predictive, not evaluative. **a)** Action potential firing (top) and lick (bottom) rates aligned to the first contact of the tongue to the rewarded lick port for the average of the 10 (out of 22) mDA neuron recordings with significant reward responses early in training (first three sessions) **b)** (top) Mean raw signal from lick port piezo strip during the recordings of each of the 10 neurons represented in (a), aligned to first rewarded lick. (bottom) Raw data rectified and convolved with a square wave to accentuate lick signals. **c)** Firing rate for mDA neurons in early (sessions 1-3), middle (sessions 4-8), and late (sessions 9+) training warped into the phase from reward delivery to first rewarded lick. **d)** Data from additional mice trained with a silenced solenoid controlling water delivery to eliminate audible noise marking reward delivery. Graph summarizes the 5 (out of 10 total) significantly modulated SNc DA neurons recorded during the first 200 trials of training.

As we noted above, overt behavior is considered an observable correlate of a subjective expectation of reward that develops during learning. The appearance of an external stimulus such as a drop of sweetened water can spur an animal to purposive action, but purposive action can occur in the absence of such stimuli as well. This raises the question of whether reward expectation that accompanies movements made in the absence of an external stimulus might likewise be associated with mDA neuron activity. For example, phasic mDA modulation has been observed around the onset of locomotion in untrained mice^16,17^, purposive actions by rodents^18–20^, and reaching movements in trained primates^21^. In our task, self-initiated movements occurred within the intertrial intervals in bouts (bout length: 3.9 ± 1.5 s) separated by periods of stillness (interbout interval: 6.4 ± 1.9 s) and 57 ± 5% of self-initiated movement bouts were accompanied by licking (Fig. 1). We observed a range of significant modulations of mDA activity at self-initiated movement onset: SNc DA neurons were more likely to be excited at movement initiation than VTA DA neurons (SNc: 19 of 66, VTA: 4 of 30), while inhibition at movement initiation was prominent in both subpopulations (SNC: 34 of 66, VTA: 17 of 30) (Fig. 4).

**Fig. 4.**
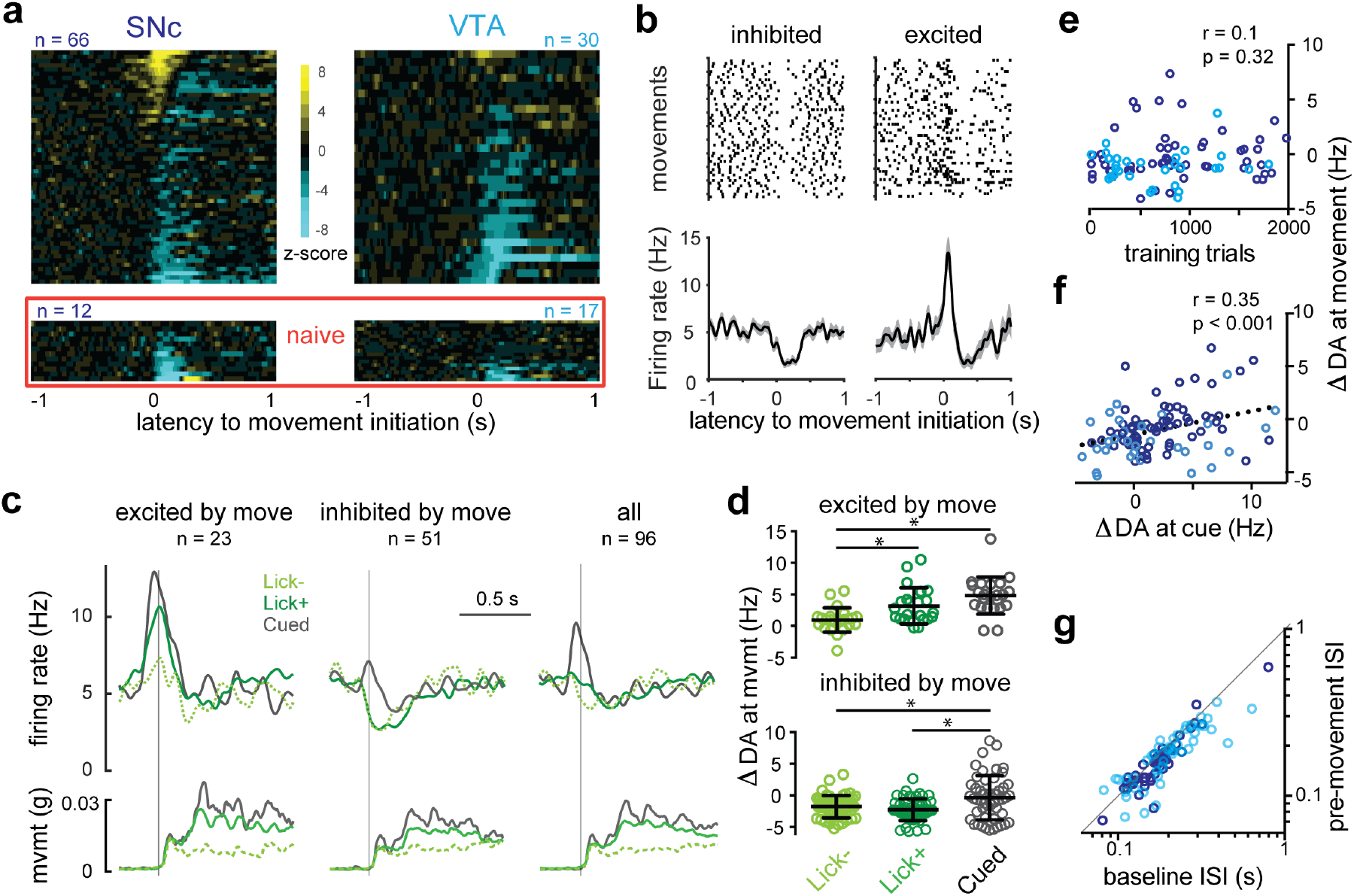
mDA movement encoding reflects reward expectation. **a)** Heat maps of z-scored PETHs for the population of SNc (left) and VTA (right) mDA neurons aligned to movement initiation either recorded within the context of the conditioning task (upper) or in animals that had yet to experience any rewards in the head-fixed context (naïve, lower). **b)** Single example mDA neurons with clear negative (left) and positive (right) modulation of firing around movement onset. Upper panels show raster plots of individual trials and lower panels show PETHs. Shaded area indicates the s.e.m. **c)** Movement-aligned PETHs from the combined population of SNc and VTA mDA neuron (n=96; right, “all”) and sorted into populations that showed significant positive modulation in the PETH (left) and negative modulation (middle). Each set of PETHs is separated into self-initiated movements absent licking (dashed green, Lick-), self-initiated movements accompanied by licking (green, Lick+), and cued movements immediately following the predictive tone (grey, Cued). Corresponding PETHs of the basket movement signal (mvmt) are shown in lower row. **d)** Quantification of individual neurons represented in C. **e)** Mean modulation of mDA neuron activity during movement initiation (ΔDA at movement) is not correlated to training level, but is correlated to mean modulation of mDA activity to tone delivery (ΔDA at cue; (**f**)). SNc (purple) and VTA (cyan) mDA neurons identified by color. **g)** Evidence of covert excitation observed by plotting the interspike interval (ISI) immediately preceding movement onset (pre-movement) against randomly selected ISIs from recording epochs without movement or cues (baseline).

These data are consistent with the proposal that phasic mDA excitation can encode parameters of movement as a necessary signal for movement initiation^16^. However, they are also consistent with the wealth of observations implicating phasic mDA neuron activity in signaling reward expectation^4^—we show above that subjective reward signals are intimately tied with the initiation of appetitive movements (Fig. 2a, e). We propose the following criteria to disambiguate these possibilities: if phasic mDA neuron excitation is necessary for movement then it should: (1) be present during all significant increases in movement amplitude, not only at transitions from rest to movement, (2) be independent of reward context, not scaled by expectation of reward, (3) be independent of reward expectation, not correlated with external sources of reward expectation, and (4) represent movement parameters within distinct subpopulations of cells^16^ not present generally in mDA neurons^12,13^. We consider each alternative in turn.

We first note that there was a clear single peak of mDA activity aligned to the onset of body movements even though the movement bouts themselves have complex profiles with multiple peaks (Fig. 4a-c). Importantly, firing rate variability over the population increased during movement initiation but not the remainder of movement bouts (COV, rest: 1.11 ± 0.04; 500-ms window surrounding initiation: 1.38 ± 0.05, p < 0.0001; remainder of movement: 1.20 ± 0.05, p = 0.08, n = 96). When we examined the largest body movements within ongoing bouts of movement we found no associated modulations of mDA neuron activity (Supplementary Fig. 2). Thus, mDA neuron activity was only modulated around the onset of movements – both for body movements and as described above for bouts of consummatory licking (Fig. 2).

We next asked whether the phasic activity of mDA neurons around the onset of movement scaled with expectation of reward. The clearest distinction in expectation is presumably between a context without any possibility of reward and one in which rewards are available. We performed additional recordings in naïve mice that had been adapted to the head-fixed behavioral chamber but never received rewards in that context. In these naïve mice, we observed no cells with significant movement-related excitation, while 10 of 12 SNc neurons and 9 of 17 VTA neurons exhibited significant inhibition (Fig. 4a, lower; Supplementary Fig. 3). Thus, mDA populations were significantly inhibited during movement initiation in the absence of rewards^17^, while clear movement-related excitation only became apparent in a reward context.

Once rewards are available behavioral correlates provide overt evidence of the magnitude of reward expectation. Anticipatory licking increases in proportion to the probability and magnitude of delayed rewards in trained animals^9,22^ and as we confirm above increases monotonically as learning progresses (Fig. 2b). Thus, self-initiated movements accompanied by licking (Lick+) represent moments of enhanced reward expectation relative to movement initiations in the absence of licking (Lick-). This allowed us to examine whether the magnitude of movement-related mDA neuron excitation scaled with changes in the magnitude of reward expectation. Indeed, in mDA neurons positively modulated by movement initiation Lick+ movements were accompanied by greater excitation than Lick-movements (3.2 ± 0.6 vs 1.0 ± 0.4 Hz, p = 0.02, n = 23), whereas inhibition in negatively modulated mDA neurons was independent of licking (−2.3 ± 0.3 vs −1.8 ± 0.3 Hz, p = 0.6, n = 51; Fig. 4c-d). Finally, we considered that movements initiated in response to a reward-predictive stimulus should be accompanied by a higher relative expectation of reward than self-initiated movements in the intertrial interval. Thus, we examined within-trial data and found that modulation of mDA neuron activity was higher for movements initiated in response to the reward-predictive tone than self-initiated movements (positively modulated: 4.8 ± 0.6 Hz, p < 0.0001 vs Lick-, p = 0.06 vs Lick+, n = 23; negatively modulated: −0.4 ± 0.5 Hz, p = 0.01 vs Lick-, p = 0.0004 vs Lick+, n = 51; Fig. 4c-d). Tone-cued excitation was noticeably blunted in neurons with net negative spontaneous movement signals (Fig. 4c), further demonstrating that movement-related inhibition is independent of—and can thus modulate—reward signaling. As noted above, our data suggest that mDA neuron responses to the reward predictive tone and reward delivery appear to be independent, but scaled by a common cell autonomous factor (Fig. 2d; Supplementary Fig. 1). If excitation around the initiation of movement reflects reward expectation then it should resemble reward signaling: it should be constant throughout training and positively correlated with the response to the tone. Indeed, the modulation of mDA neuron activity at movement initiation was not correlated with training extent (p = 0.32, Fig. 4e, n = 96) but it was positively correlated with the magnitude of the tone response in individual neurons (r= 0.35, p < 0.001, n = 96, Fig. 4f).

Finally, we explore whether our data support the view of a parallel architecture in which distinct mDA subpopulations encode movement or reward information. SNc and VTA DA neurons both predominantly responded to rewards and reward-predictive cues (Fig. 2), as observed once previously in rodents^23^, arguing against a simple anatomical division of labor. Furthermore, neurons lacking excitation to reward cues were also likely to lack excitation at self-initiated movement (Fig. 4f), and only 1 of 21 mDA neurons that lacked significant reward responses exhibited significant excitation at movement initiation. Excitation at reward delivery and movement initiation are thus clearly not mutually exclusive. We next asked whether neurons that lacked significant movement-related excitation reflected categorical differences in afferent connectivity or graded differences in the strength of input. We closely examined activity around movement initiation in these neurons by attempting to infer the presence of excitation as a change in the interspike interval (ISI) just before movement-related pauses (see Methods and Supplementary Fig. 3E). This analysis uncovered significant excitation in the form of a spike phase advance (9 ± 2% faster, p < 0.001, n = 73, Fig. 4g) that was “covert” in the sense that it was not apparent from averaged peri-event time histograms. These analyses argue against categorical restriction of putative movement-encoding afferents to defined subpopulations of mDA neurons^16,17^. However, SNc neurons are clearly more sensitive to excitation around self-initiated movements than VTA neurons, possibly reflecting graded biases in afferent input across the two populations^24^. It is possible that this bias coupled with a threshold for detection of activity (*e.g*. calcium imaging) could produce apparent categorical differences.

Taken together these results strongly support a model in which the phasic responses of mDA neurons to reward predictive stimuli and initiations of purposive movements reflect a common correlate: reward expectation. Furthermore, there are multiple sources of reward expectation information that can change independently (Fig. 2) and appear to be additive (Fig. 4). On the one hand, these data replicate recent observations of movement-related mDA activity in naïve animals while also reconciling them with prior results that activity is primarily related to reward predictive stimuli^16,17,21,25^. On the other hand, these results conflict with the commonly held view that the computation underlying phasic modulation of mDA activity is the subtraction of inhibitory prediction signals from excitatory actual reward signals (Reward Prediction Error, RPE^5^). First, the magnitude of the mDA response is determined by the animal’s behavior rather than being a product of external stimuli because it depends upon the time at which appetitive movements are initiated. Second, mDA neuron activity does not reflect the evaluation of a received reward but rather an expectation associated with the action to collect the reward. This latter point argues fundamentally against a computation of “error”, because there is not an evaluative component of mDA neuron activity even very early in training (Fig. 3). However, as we will show, these two principles of mDA neuron activity are in fact sufficient to produce RPE *correlates* in mDA neuron activity (which we replicate here), while challenging the computation proposed to underlie those correlates.

To examine RPE correlates in mDA neuron activity we omitted the reward predictive tone on 30% of trials in select recording sessions. This produced ‘unpredicted’ water delivery events that we could compare to the predicted water delivery considered to this point. To provide behavioral evidence that the tone predict water delivery we examined how rapidly mice initiated a consummatory bout of licking following reward delivery across training. By the middle training epoch (sessions 4-8), a predictive tone indeed reduced the time to initiate consumption after reward delivery (Fig. 5a; Supplementary Fig. 4).

**Fig. 5.**
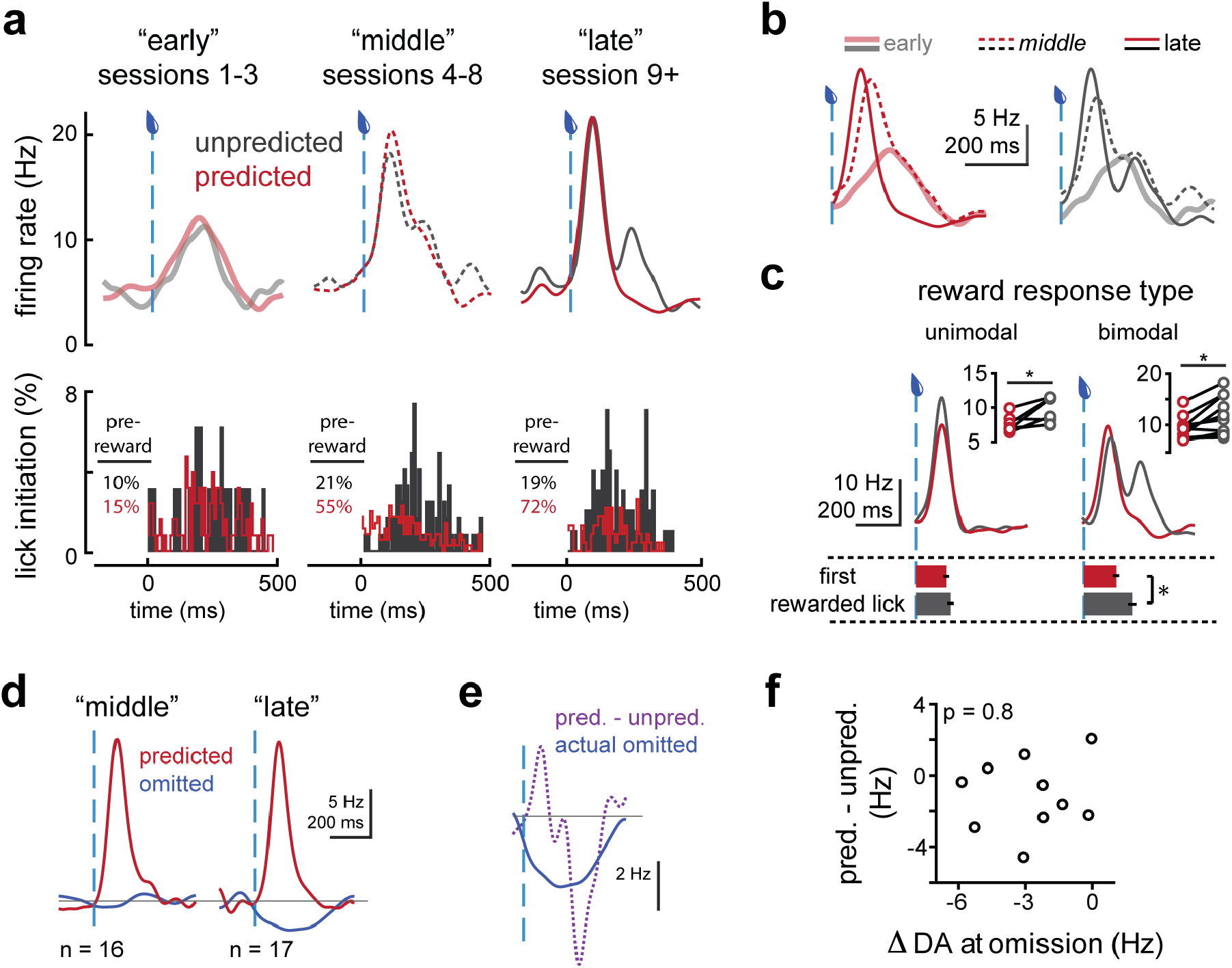
RPE correlates in mDA neurons reflect the presence or absence of action initiation at the US. **a)** PETHs of mDA activity (upper) and distribution of lick bout initiation times (lower) aligned to predicted (red) or unpredicted (dark grey) reward delivery for early (left), middle (center), and late (right) training sessions. The percent of trials with a previously initiated lick bout ongoing during reward delivery (“pre-reward”) are inset. **b)** PETHs from A overlaid and sorted by predicted (left, red) and unpredicted (right, black) reward delivery. **c)** PETHs from significantly-modulated mDA neurons in late sessions divided into short and long first rewarded lick intervals for predicted (red) and unpredicted (dark grey) reward delivery. Lower row shows the mean and s.e.m. of the latency to the first rewarded lick for each population **d)** Comparison of PETHs aligned to reward delivery in predicted trials (red) and to the moment of omitted reward delivery in trials with no reward delivery following the predictive tone (blue) are shown for middle (left) and late (right) training epochs. **e)** PETHs aligned to reward delivery for actual omission trials (blue) compared with the inferred, putative subtractive prediction effect (pred. PETH – unpred. PETH, purple). **f)** Inferred effect of prediction on mDA modulation by reward delivery (pred.-unpred.) plotted as a function of the mean modulation a movement initiation (F). Inset p-value indicates the result of significance testing of the correlation between the plotted variables.

We next asked how mDA activity at the time of reward delivery was modulated by the presence or absence of a predictive stimulus. In early training, mDA responses were relatively small and slow, and unaffected by prediction (pred. vs unpred.: 2.3 ± 1.6 vs 1.5 ± 1.2 Hz, p=0.4, n=12, Fig. 5a). mDA reward responses remained unaffected by prediction in middle training (pred. vs unpred.: 4.3 ± 0.6 vs 4.1 ± 0.7 Hz, p = 0.65, n = 30) despite evidence of learning in the form of behavioral responses to the predictive tone (Fig. 2a) and a more rapid reaction to water delivery (Supplementary Fig. 4). It was not until late training that unpredicted rewards began to evoke relatively larger mDA reward responses (3.1 ± 0.6 vs 4.5 ± 1.0 Hz, p = 0.03, n = 24), with the difference apparent in the delayed component of a bimodal response. The eventual emergence of positive RPEs here is consistent with previous observations in well-trained mice^22^, but it has not heretofore been appreciated the degree to which the RPE correlate lags behavioral evidence of learning.

Why does it require so much training for prediction to have an effect on mDA reward responses? We propose that prediction only indirectly modulates mDA activity by changing the structure of behavioral responding to rewards (Supplementary Fig. 5). This is supported by the following observations. The delayed component of the mDA reward response had a very similar latency and magnitude to the reward response early in training (Fig. 5b). Moreover, the time of the second peak of the mDA reward response corresponded well to a second mode in the distribution of lick bout initiations following reward delivery (Fig. 5a, right). A minority of mDA neurons exhibited only a unimodal reward response, yet still encoded a difference between predicted and unpredicted rewards (Fig. 5c left, pred. vs unpred.: 7.6 ± 0.5 vs 9.4 ± 0.7, p=0.03, n= 7). This indicated that a delay was not necessary for positive RPEs to be signaled. Importantly, during the recording of these mDA neurons’ activity the latency to reward collection was similar between predicted and unpredicted trials (pred vs unpred: 142 ± 14 vs 162 ± 17 ms, n = 7, p = 0.06). By contrast, reward collection was significantly delayed (~50 ms) in unpredicted trials during recordings in which individual mDA neurons displayed bimodal reward responses (pred vs unpred: 155 ± 9 vs 213 ± 27 ms, n = 11, p = 0.02). Thus, the second mode of mDA neuron activity observed in unpredicted trials corresponds to the latency of the appetitive action initiated by reward delivery (*i.e*. a consummatory lick bout).

Bimodal mDA reward responses have been observed across species and have been proposed to reflect an initial salience signal that is insensitive to prediction followed by a delayed evaluation signal sensitive to prediction^4^. Instead, early in training (Fig. 5b) we observed that a unimodal, relatively delayed signal corresponded to unpredicted reward collection (Fig. 3), and a larger, faster component emerged as training progressed, corresponding to the learning of faster behavioral responses (Supplementary Fig. 4) to the sensory components of reward delivery. In late training, the delayed mode of the mDA reward response correlates with delayed initiation of appetitive action specific to unpredicted reward delivery. Thus, a model in which mDA activity reflects the summation of multiple sources of reward expectation can account for the positive RPE correlate as well as providing an explanation for why it only emerges once learning is essentially complete (Supplementary Fig. 5).

We finally explored mDA neuron activity during trials in which predicted reward delivery is omitted – a key piece of evidence that mDA activity computes and reports RPEs (Supplementary Fig. 5). Brief pauses in mDA neuron activity during omitted rewards have been argued to provide direct evidence that a ‘negative prediction’ accounts for the relative suppression of mDA neuron responses to predicted reward delivery^4,26,27^. Our data reveals inhibition of mDA neurons during movement initiation even in naïve animals (Fig. 4; Supplementary Fig. 3). Is it possible that inhibition upon reward omission results from behavior that emerges specifically in omission trials and thus is distinct from the mDA response to predicted reward delivery? To examine this possibility, we omitted rewards on a subset (<20 %) of trials during select recordings in middle and late training. We only observed a significant pause in mDA activity to omitted reward delivery late in training (middle training: 5.4 ± 0.7 to 5.7 ± 0.9, p = 0.73, n = 16; late training: 5.3 ± 0.6 to 2.6 ± 0.4, p <0.0001, n = 17; Fig. 5d). This is consistent with the late emergence of positive RPE correlates, but once again substantially lags behavioral evidence of learning and mDA encoding of a reward predictive stimulus. Importantly, if omission-related pausing reveals an inhibitory signal (negative prediction) inferred to be present during predicted rewards, the actual inhibition and the inferred inhibition should be well-matched since reward omission trials are randomly interleaved. In contrast to this prediction, the suppression of mDA neuron activity on interleaved omission trials was not correlated with either the timing (Fig. 5e) or the magnitude (Fig. 5f) of the ‘negative prediction’ inferred from rewarded trials. Together, these data argue that positive and negative components of the RPE correlate are mediated by dissociable mechanisms (Supplementary Fig. 5).

Here we show that the positive, phasic responses of mDA neurons reflect the summation of multiple reward expectation signals, but not the evaluation of collected reward. These moments of reward expectation are driven by external or internal stimuli, and are closely associated with the self-initiation of purposive movements. As a consequence, apparent RPE correlates in mDA neuron activity, at least in naïve mice being introduced to an associative trace conditioning paradigm, do not, indeed cannot, report errors in prediction. Rather, mDA neuron activity reflects the changing summation of external (specified by the experimenter/environment) and internal (controlled by the animal) causes of expectation that is a function of learning (Supplementary Fig. 5). We suggest that these distinctions were only apparent by examining identified mDA neurons during novel learning.

Our model has several important implications for the understanding of how multiple learning circuits interact to mediate learned behavior and the mechanistic role phasic mDA neuron activity may play in that process. For example, here we replicate the RPE correlate of mDA neuron activity, but show that it is explained by a computation that does not invoke errors. This eliminates the explanatory gap filled by a proposed negative prediction signal. While future studies will be required to extend these observations to highly-trained animals and other species, we note that it has been emphasized that in trained animals unambiguous negative prediction signals have been difficult to identify despite significant effort^5^. This also has important conceptual implications. Our data demonstrate that mDA neuron activity does not reflect a delayed “evaluation” phase of reward responses required to establish the value of an “actual” reward. In naïve animals mDA activity anticipates reward collection (Fig. 3)^7^. Thus, mDA neuron activity, at least during early learning, is not readily reconciled with changes in a scalar expected value function^28–30^. While these observations might be reconciled with the addition of post hoc conjectures to an RPE computation, a simple model with substantial explanatory power is that mDA neuron activity reflects the increasingly precise coordination of external, sensory predictors and reliable behavioral responses to those predictors during initial learning. Our model both produces RPE correlates and makes several novel predictions confirmed here (Supplementary Fig. 5).

Our data are consistent with the fundamental insight that mDA neuron activity reflects subjective reward expectation rather than a necessary component of motor pathways^21,31^. Indeed, these observations during novel learning provide strong evidence that mDA activity may *only* reflect changes in reward expectation rather than also reflecting differences between expectation and evaluation of the actual reward received or objective parameters of movements. In this way, RPE correlates emerge following behavioral adaptations to learning due to the precise timing of external and internal causes of reward expectation. This is consistent with causal evidence that exogenous induction of RPE correlates in mDA neuron activity is sufficient for learning^32–34^. However, this mechanistic revision is fundamental because error-based models were first articulated to explain novel learning and mDA neuron activity is known to be necessary for novel learning. While error signals can, unquestionably, be useful to drive learning^2,11^ we note that Hebbian-like learning rules based upon correlations between input (external) and output (internal) are sufficient for learning and can be equivalent to error-based rules^35^. These arguments suggest that a Hebbian-like teaching signal in the activity of mDA neurons, allowing a novice animal to learn from its successes rather than its errors, may be sufficient for the initial emergence of adaptive, reward-seeking behavior.

## Methods

### Animals

All procedures and animal handling were performed in strict accordance with protocols (11-39) that were approved by the Institutional Animal Care and Use Committee (IACUC) and consistent with the standards set forth by the Association for Assessment and Accreditation of Laboratory Animal Care (AALAC). For behavior and juxtacellular recordings we used 11 adult mice (10-24 weeks old) resulting from the cross of DAT^IRES*cre*^ (The Jackson Laboratory stock 006660) and Ai32 (The Jackson Laboratory stock 012569) lines of mice, such that a Chr2/EYFP fusion protein was expressed under control of the dopamine transporter Slc6a3 promoter to specifically label dopaminergic neurons. All animals were handled in accordance with guidelines approved by the Institutional Animal Care and Use Committee of Janelia Research Campus. Animals were housed on a 12-hour dark/light cycle (8am-8pm) and recording sessions were all done between 9am-1pm. Following at least 4 days recovery from headcap implantation surgery, animals’ water consumption was restricted to 1 mL per day for at least 3 days before training. Mice underwent daily health checks, and water restriction was eased if mice fell below 75% of their original body weight. Mice were habituated to head fixation in a separate area from the recording rig in multiple sessions of increasing length over 3 days, including manual water administration through a syringe. Mice were then habituated to the recording rig for at least two 30+ minute sessions before recordings were attempted or training commenced.

### Behavioral training

Mice head-fixed while resting in a spring-suspended basket were trained to learn a classical (Pavlovian) auditory trace-conditioning paradigm. The reward consisted of 3 μL of water sweetened with the non-caloric sweetener acesulfame potassium delivered through a lick port under control of a solenoid. In the first training session, a 0.5 s, 10 kHz tone preceded reward delivery by 1.5 s on 100% of trials. In subsequent training sessions, “unpredicted” reward responding was assessed by randomly omitting the predictive tone on 30% of trials during select blocks of trials, with the result that the proportion of unpredicted reward trials never exceeded 10% over any given training session. “Omitted” reward responding was assessed by randomly omitting the reward following a predictive tone on 30% of trials during select blocks of trials, such that the proportion of omitted reward trials never exceeded 10% over any given training session. Intertrial intervals were chosen from randomly permuted exponential distributions (means of ~10, 25, or 50 s) every 20 trials in order to fully disrupt reliable estimation of intertrial interval while keeping the mean interval tractable for recording. Ambient room noise was 50-55 dB, while an audible click of ~53 dB attended solenoid opening upon water delivery and the predictive tone was ~65 dB loud. It should be noted that these stimuli are all quieter than the 72-90 dB stimuli that were reported to activate mDA neurons via their salience in primates^12^. Indeed, in naïve animals no significant modulation was apparent to the tone (Fig. 2a, left), and the timecourse of modulation by reward delivery early in training (Fig. 5a,b) was not consistent with proposed salience signaling in mDA neurons. However, to control for such signaling as an alternative explanation for the predictive nature of the mDA reward response, we recorded from additional mice trained with an inaudible solenoid (The Lee Company, LHQA1231220H).

### Behavioral and electrophysiological measurements

Individual licks were timestapped according to deflections of a piezo strip supporting the lick port. Piezo signals were high-pass filtered at 0.1 Hz, then rectified and smoothed by convolving with a square wave in order to facilitate identification of individual tongue strikes during high frequency lick bouts.

Basket movements were recorded by a triple-axis accelerometer (Adafruit, ADXL335) attached to the underside of a custom-designed 3D-printed basket suspended from springs (Century Spring Corp, ZZ3-36) well suited to allow robust movements but fully support the ~20-25 g body weight of adult mice. Basket height relative to the point of head fixation was set so that a mouse’s back was at a ~30° angle to the plane of the its headcap. This positioning was comfortable for the animals and minimized the translation of body movement to movement of the brain with respect to the skull, allowing for more stable juxtacellular recordings. Raw accelerometer signals summed across all axes were used to identify transitions from rest to movement (movement initiations). Relative basket position was tracked by low-pass filtering accelerometer data at 2.5 Hz to enrich for the signal corresponding to forces due to the angle of the accelerometer with respect to the earth. Both lick and movement events were detected by threshold crossing but were timestamped according to the their earliest deviation from the previous baseline signal.

Electrophysiological recordings were made with a Multiclamp 700B amplifier (Molecular Devices) interfaced to a computer by an analog-to-digital converter (National Instruments, PCI 6259) controlled by Axograph X recording software (www.axograph.com). Spikes were recorded in current clamp mode, AC-coupled at 1 Hz, then high-pass filtered at 300 Hz to facilitate simple threshold crossing analysis to generate spike time stamps. Data were smoothed by convolving with a Gaussian function with a 20 ms decay.

The above signals as well as the command signals that spanned the predictive tone length, the reward solenoid open time, and the on time of the laser used for optotagging were synchronously recorded and digitized (at 1 kHz for behavioral data, 30 kHz for electrophysiology data) with a Cerebus Signal Processor (Blackrock Microsystems). Stimulations and cue deliveries were coordinated with custom-written software using Arduino Mega hardware (www.arduino.cc). Data was analyzed using Matlab software (Mathworks).

### Juxtacellular recording

A small craniotomy (<200 μm diameter) was made over the recording site (from bregma: −3.2 posterior, 0.5 lateral for VTA, −3.0 posterior, 1.5 lateral for SNc) at least 4 hours prior to recording. Exposed brain tissue was kept moist with phosphate-buffered saline at all times, and craniotomy sites were covered with Kwik-Sil elastomer (WPI) outside of the recording session. Borosillicate glass pipettes (Sutter, BF165-120-10) were pulled to a long taper (~10 mm taper) with a ~1-2 μm tip (resistance 4-14 mOhm) with a P-97 micropipette puller (Sutter). Pipettes were filled with 0.5 M NaCl solution and mounted in a holder with a side port (Warner, PE30W-T17P) to allow insertion of a fiber (105 μm core, 0.22 NA, Thorlabs) that was coupled to a 473 nm laser (50mW, OEM Laser Systems) to carry light to the pipette tip. An AgCl pellet reference electrode was placed in the well of saline that covered the craniotomy site.

Pipettes were lowered through the brain with a micromanipulator (Luigs and Neumann) while a small cycling current injection allowed monitoring of resistance changes across the pipette tip. In addition to stereotactic coordinates, dopaminergic regions could be further targeted empirically by monitoring extracellular potentials in response to brief (<5 ms) flashes of blue light through the pipette tip (Fig. 1E). The amplitude of population responses closely corresponded to patterns of dopaminergic cell body enrichments predicted from standard coordinates^8^. Deviations from that correspondence represented errors introduced through individual anatomical variation or the plane of headcap attachment to the skull, and referencing of the population response allowed correction on subsequent passes, greatly increasing the yield of identified mDA recordings across the life of each animal. Once within a dopaminergic region, the pipette tip was advanced by 1-2 μm steps until a steep increase in resistance was detected. The pipette was then advanced 5-10 μm until positive-going spikes were resolved well above noise (>~0.5 mV). ChR2-expression was assayed with 0.5 s 473 nm light stimulations, either in bursts of 1 ms flashes at 10 Hz or continuous pulses. Cells were not re-identified for the remainder of the recording session so as not to interfere with physiological responses. Cells were held for a median of 11 trials (interquartile range 7:19) and a median of 27 self-initiated movements (IQR 14:47). In some pilot experiments, identified mDA cells were labeled in the juxtacellular configuration for future recovery by including Neurobiotin (Life Technologies) in the pipette solution and entraining spiking with 2 Hz, 50% duty cycle current injections of the minimum amplitude required to entrain spiking (>1 nA). During pilot experiments, 0 of 20 neurons in the first 2 mm of tissue overlying the targeted areas were significantly modulated by stimulation with max laser power (~26 mW out of the fiber tip), indicating that activation due to heating or other nonspecific processes was not a concern.

We quantified tone and reward responses by averaging firing rates over a 500-ms window following the cue and subtracting the average baseline firing rate in the 1000 ms preceding tone delivery^22^. We quantified movement responses by averaging firing rates over a 300-ms window beginning 50 ms before movement initiation and subtracting baseline firing rates—this window was chosen in order to capture the bulk of both excitatory and inhibitory responses across different neurons. Slightly different windows were used for one-tailed tests of significant modulations of activity by movement initiation in single cells: for excitation, a 200 ms window beginning 100 ms before initiation was used; for inhibition, a 300 ms window beginning at the moment of movement initiation was used. These parameters better described the activity during the optimal windows for each signal.

### Statistical analysis

Paired comparisons within individual cells were made using the Students’ t-test. Multiple comparisons were made using a one-way ANOVA with Tukey’s post-hoc multiple comparisons test (Graphpad Prism). Errors are reported as standard errors of the mean (s.e.m.).

### Histology

To recover locations of recorded neurons in pilot experiments, mice were killed by anesthetic overdose (isoflurane, >3%) and perfused with ice-cold phosphate-buffered saline (PBS), followed by paraformaldehyde (4% wt/vol in PBS). Brains were post-fixed for 2 h at 4° C and then rinsed in saline. Whole brains were then sectioned (100 μm thickness) using a vibrating microtome (VT-1200, Leica Microsystems) and incubated with Alexafluor 594-conjugated streptavidin was used to visualize Neurobiotin-labeled neurons against the backdrop of EYFP-expressing mDA neurons.

---

**Supplementary Figure 1.**
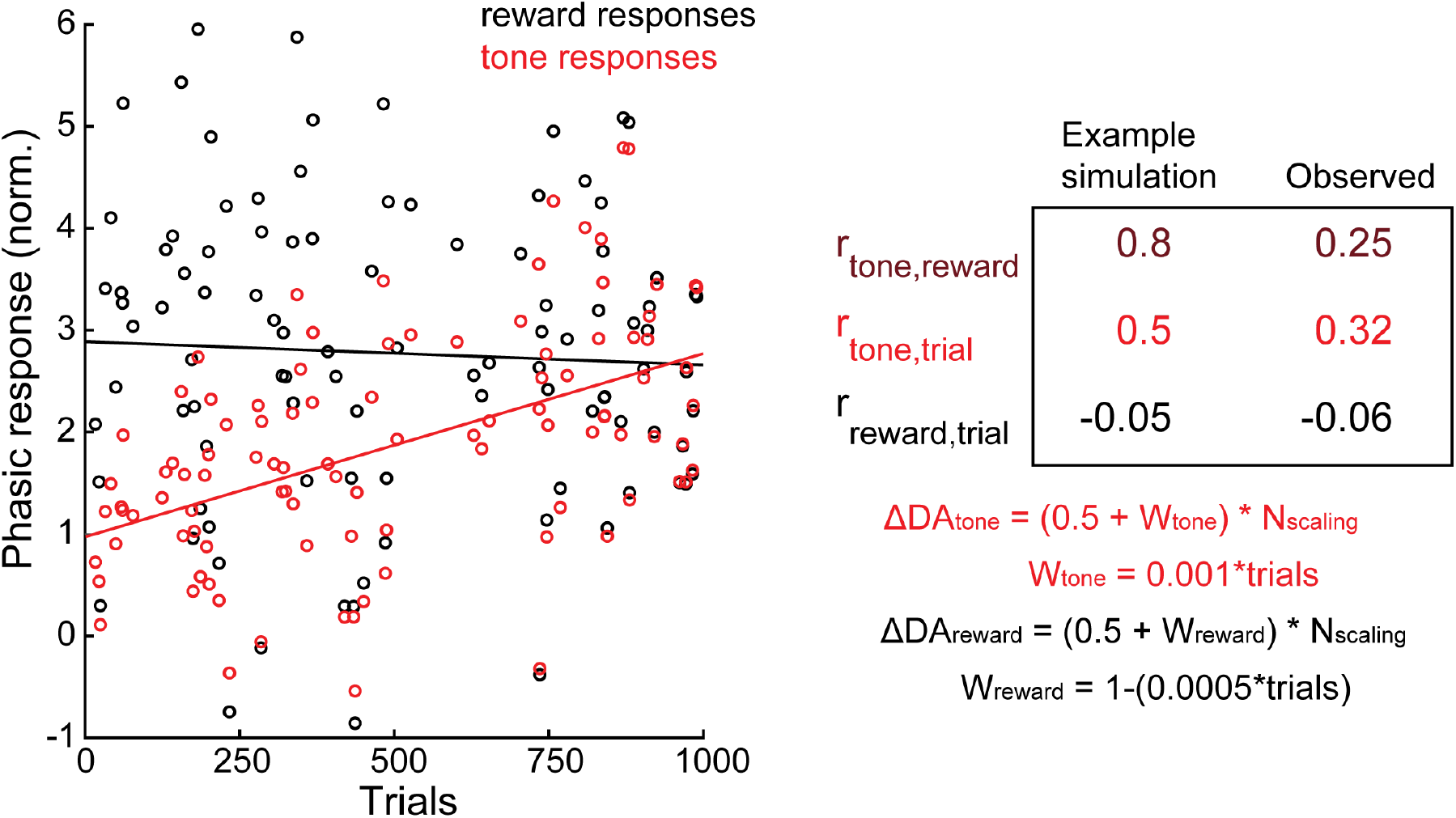
A modal of cue and reward activity incorporating independently-learned responses scaled by a common factor replicates observed relationships in the data. (left) Example results plotting the phasic modulation of activity (arbitrary units) as a function of trials for a simulation of the equations governing the change in dopamine neurons activity (ΔDA) at the time of the predictive tone (ΔDA_tone_) and the reward (ΔDA_reward_). Equation is explicitly shown at right. Random numbers were drawn from a normal distribution (N_scaling_). (right) Correlation coefficients were calculated for all trials in the simulation (N=1000) for comparison with ‘Observed’ correlations taken from the main text. Throughout the figure red corresponds to the tone responses and black corresponds to reward responses.

**Supplementary Figure 2.**
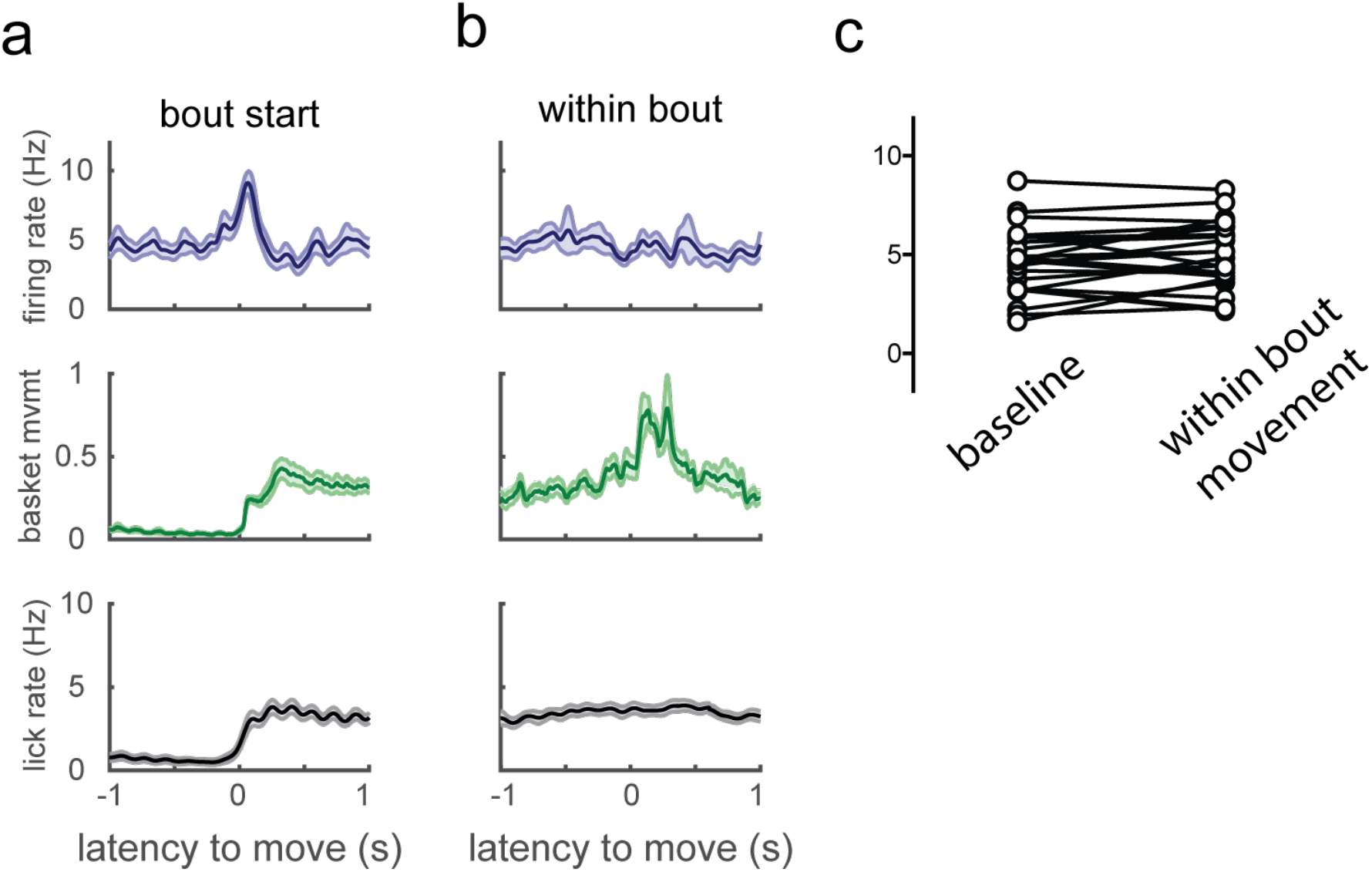
mDA neurons do not encode within-bout movement. **a)** Average firing rate (top), basket movement (middle), and lick rate (bottom) for 23 mDA cells significantly excited at movement initiation. **b)** Same as (a), but for the point of maximum basket displacement within in each movement bout excluding the first 500 ms surrounding movement initiation. **c)** No significant difference was observed between baseline rates and rates during the within-bout movements shown in B (p = 0.6).

**Supplementary Figure 3.**
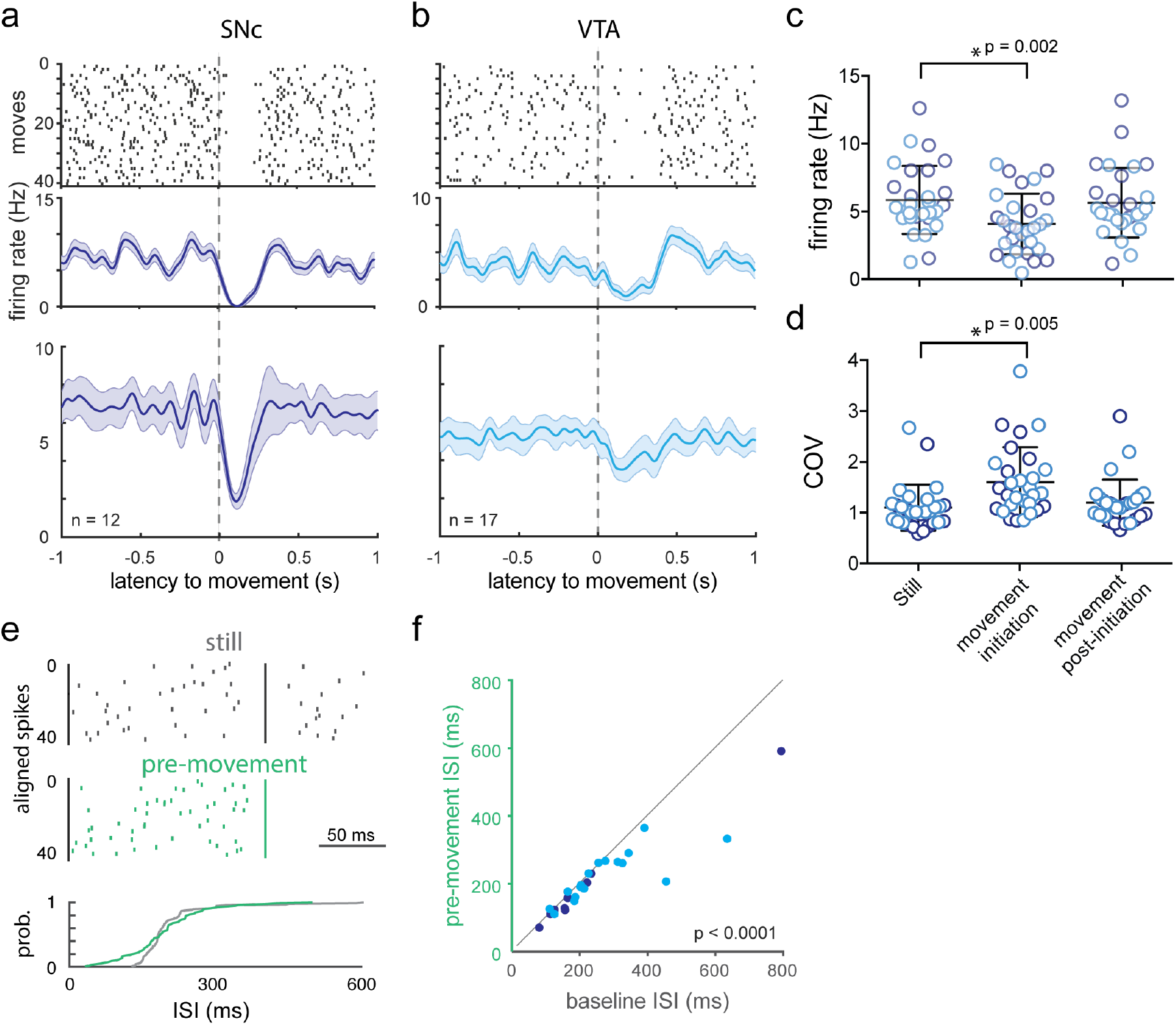
mDA neurons are predominantly inhibited at movement initiation in the absence of a reward context. **a)** Raster plot (top) and PETH (middle) aligned to movement initiation for example SNc DA neuron activity and for the population (bottom), recorded in naive mice which had received no rewards in the training rig. **b)** Same as (a), but for VTA neurons. Average firing rates (**c**) and coefficients of variation (**d**) were calculated for periods of stillness, a 500 ms window beginning at movement initiation, and the remainder of movement bouts following the initiation period. **e)** Comparison of spike rasters aligned to random spikes drawn from periods of stillness (top) vs aligned to the last spike before the movement initiation-related pause (middle), with the cumulative density function at bottom, from the cell shown in (a). **f)** Interspike intervals from the 1 sec baseline prior to movement onset were significantly slower than for the last spikes before movement onset.

**Supplementary Figure 4.**
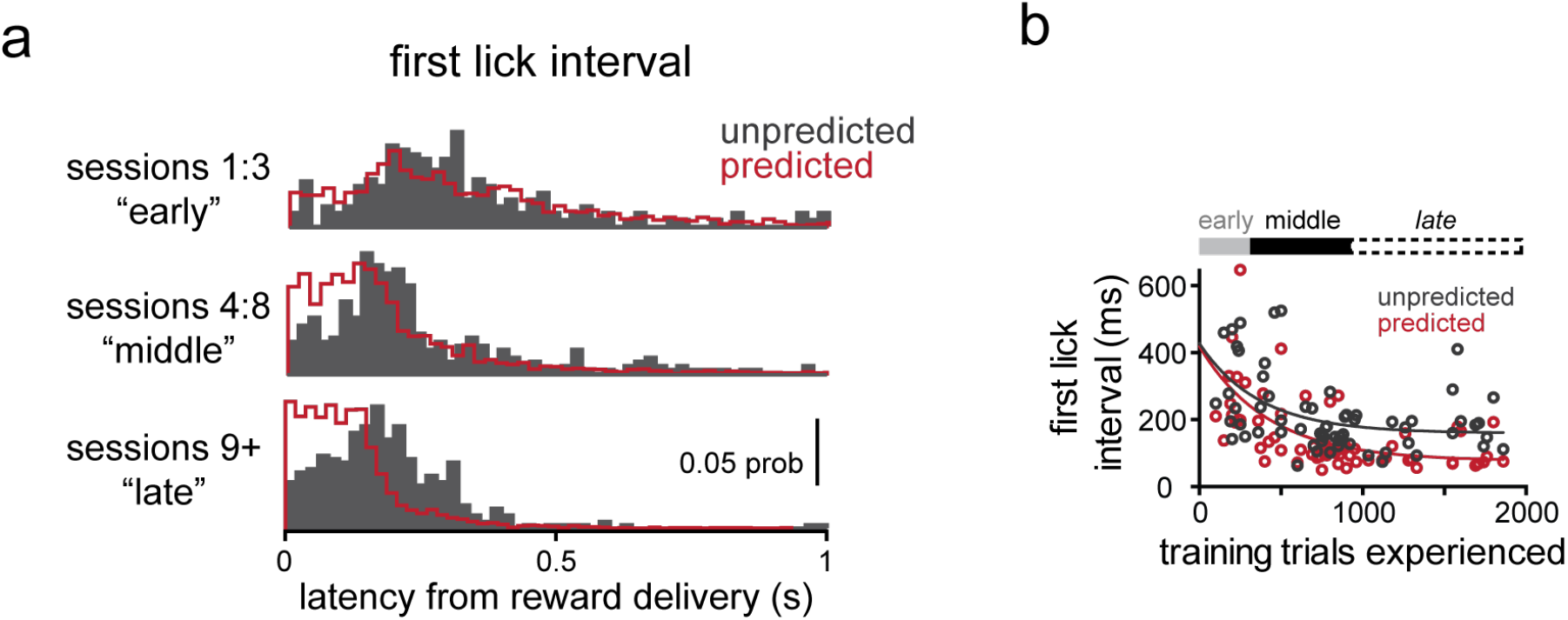
Differences in lick responding across training reflect shifts in the timing of action initiation. **a)** Timing of the first contact of the tongue to the lick port following reward delivery, in early, middle, and late training. **b)** Mean reward->first lick interval during each individual neuron recording session.

**Supplementary Figure 5.**
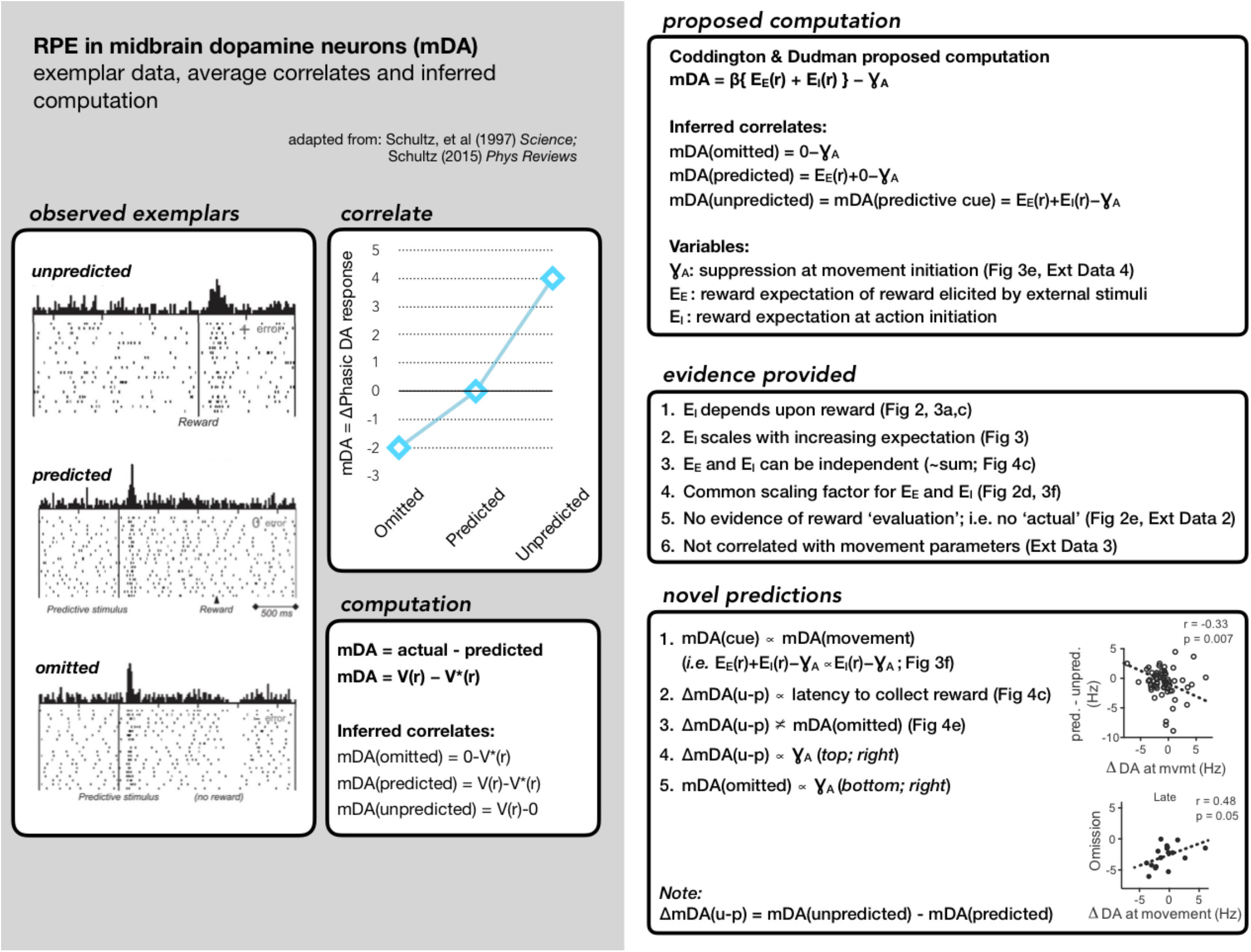
Summary of currently accepted model of Reward Prediction Error (RPE) correlates and underlying RPE computation (left, gray). Summary of the revised computation proposed here, primary evidence provided throughout this manuscript, and a set of novel predictions that follow from our proposal and are confirmed in this dataset (right column).

## Acknowledgments

*This work was supported by the Howard Hughes Medical Institute. J.T.D. is a Janelia Group Leader. Thanks to the Dudman lab, Vivek Jayaraman lab, Brett Mensh, Albert Lee, Gerry Rubin, and Jeremy Day for project feedback*.

## Author Contributions

Data collection and analysis were performed by L.T.C. with input from J.T.D. All other aspects of the work were the product of both authors.

